# An oligosaccharide analog exhibited antifungal activity by misleading cell wall organization via targeting PHR transglucosidases

**DOI:** 10.1101/2021.07.16.452596

**Authors:** Ruilian Li, Limeng Zhu, Dongdong Liu, Wenjing Wang, Chen Zhang, Siming Jiao, Jinhua Wei, Lishi Ren, Yuchen Zhang, Xun Gou, Xianghua Yuan, Zhuo A. Wang, Yuguang Du

## Abstract

The fungal cell wall is an ideal target for the design of antifungal drugs. In this study we used an analog of cell wall polymer, a highly deacetylated long-chain chitosan oligosaccharide (HCOS), to test its effect against pathogenic *Candida* strains. Results showed that HCOS was successfully incorporated into the dynamic cell wall organization process and exhibited an apparent antifungal activity against both plankton and mature fungal biofilm, by impairing the cell wall integrity. Unexpectedly, mechanistic studies suggested that HCOS exerts its activity by interfering with family members of PHR β-(1,3)-glucanosyl transferases and affecting the connection and assembly of cell wall polysaccharides. Furthermore, HCOS showed great synergistic activity with different fungicides against *Candida* cells, especially those in biofilm. These findings indicated HCOS has a great potential as an antifungal drug or drug synergist and proposed a novel antifungal strategy with structure-specific oligosaccharides mimicking cell wall polysaccharide fragments.

**IMPORTANCE:** Fungal infections have always been a puzzle in clinical medicine. Only a few antifungal drugs are available for medical usage and the widespread use of antifungal drugs increased the incidence of drug resistance. It is an urgent need for the development of novel treatment strategies against fungal infections. In this study, we proposed a novel strategy targeting to fungal cell wall against *C. albicans.* To our knowledge, it is the first study to show a cell wall polysaccharide fragment analog integrate into and interfere with the fungal cell wall, indicating a novel antifungal strategy using structure-specific oligosaccharides mimicking cell wall polysaccharide fragments.

## INTRODUCTION

Worldwide, more than 400,000 life-threatening invasive fungal infections occur yearly (1–3). Since fungi are eukaryotes like their human host, the number of targets available for antifungal drug development remains limited. The fungal cell wall is an ideal target for the design of antifungal drugs because it is composed almost exclusively of molecules that are not presented in the human body yet are essential for fungal growth virulence and viability (4). The new class antifungal compound targeting cell wall, such as echinocandin, has been widely used in medical practice (5).

The central core of fungal cell wall was composed of branched β-(1,3)-glucan cross‐linked to chitin, that forms a tight network to prevent osmotic or mechanical injuries to the cell (6, 7). β-(1,3)-glucan, the most abundant structural component of the fungal cell wall, is synthesized as a linear polymer (8), which is elongated and branched further by the glycosylhydrolase 72 (GH72) family glucanosyltransferase Gas/Gel/Phr/Epd proteins (9). Chitin, a linear polysaccharide consisted of β-1,4-linked N-acetylglucosamine residues (10), takes up 1% to 15% of the fungal cell wall mass (11). Chitin is essential for cell survival because of its significant contribution to the rigidity and strength of the cell wall (6, 8, 12). Studies showed that cell wall chitin possesses diversity in structure and function through variations in the length and degree of deacetylation of the microfibrils (13). Chitin polymers can be deacetylated by chitin deacetylases to generate chitosan, a polymer of glucosamine residues (8). The degree of deacetylation can reach up to 70-80% in some fungi, such as *Cryptococcus neoformans* (14, 15). Chitosan is structurally similar to chitin and plays an indispensable role in maintaining cell barrier functions and integrity during cell growth (16). Aberrant chitin deacetylation causes a decreased rate of growth and reduced pathogenicity (17, 18).

The cell wall is a dynamic structure that undergoes constant remodeling, including modulation of the distribution and crosslinking, essential for cell integrity and survival (19, 20). Monosaccharide analogs had been used to interfere with the cell wall glycan biogenesis or explore the glycan biosynthetic machinery by “hijacking” glycan biosynthesis pathways (21–23). It gave us a hint of a novel antifungal strategy: an analog of cell wall fragment/fiber incorporated into the process of fungal cell wall organization to impair the cell integrity. In this study, a chitosan fragment with defined structural characteristics (MW: 5 kDa; the degree of deacetylation: above 90%) was used to test this hypothesis in *Candida albicans*, the most predominant fungal species causing superficial to life-threatening systemic infections (24–26). The chitosan fragment was structurally similar to the cell wall chitosan polymers, but likely has a different degree and pattern of deacetylation. Our studies showed the fragment integrated into the cell wall and influenced the assembly process of cell wall remodeling, exhibiting excellent potential as a synergist used in antifungal therapy, especially for mature *Candida* biofilm, a resistance form of *C. albicans*.

## RESULTS

### The long-chain chitosan oligosaccharide (HCOS) exhibited apparent antifungal activity by impairing the cell wall integrity

Two chitosan oligosaccharide products, a long-chain chitosan oligosaccharide (HCOS, average molecular weight 5 kDa, degree of deacetylation > 90%) and a shorter one (LCOS, average molecular weight 0.8 kDa, degree of deacetylation > 90%), were used to assess the antifungal activity against planktonic *C. albicans* cells or its biofilms. HCOS but not LCOS showed evident inhibitory activities against fungal cells in both two lifestyles (Fig. 1A and B). It should be noted that the fungicidal activity of HCOS became more potent on the late-stage biofilm, which usually showed much greater resistance to antifungal drugs, such as caspofungin (Fig. 1C). HCOS treated cells were also more sensitive to cell wall stressors (Fig. 1D). Electron microscopic studies showed that after exposure to a fungicidal concentration of HCOS, *C. albicans* cells exhibited a porous cell surface appearance (Fig. 1E) and an ambiguous and sparse cell wall layer (Fig. 1F). These results demonstrated that HCOS affected the integrity of the cell wall structure.

**FIG 1.**
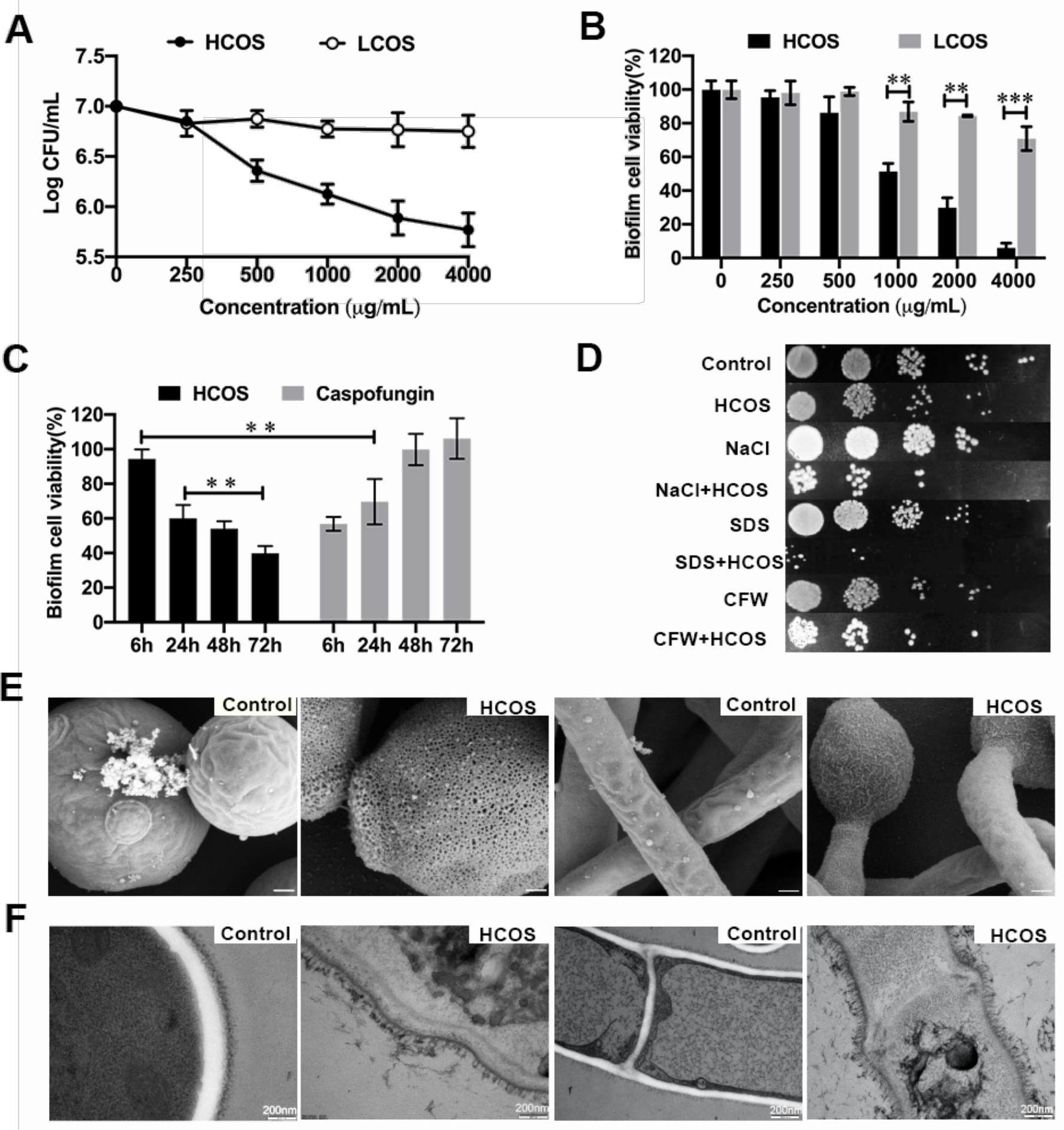
HCOS exhibited antifungal activity by impairing the cell wall integrity. (A) Planktonic *C. albicans* were treated with a series of concentrations of HCOS for 24 h and cell viability was assessed by colony counts. Data are represented as means ± SD (n=3). (B) Mature *C. albicans* biofilms were exposed to HCOS on different concentrations for 24 h, the biofilm cell viability was determined by XTT assay and normalized to the control. Data are represented as means ± SD (n=6). (C) *C. albicans* biofilms were developed in proper conditions for 6 h, 24 h, 48 h, respectively, and then exposed to HCOS (500 μg/mL) or CAS (1 μg/mL) for 24 h. The biofilm cell viability was determined by XTT assay. Data are represented as means ± SD (n=6) of cell viability compared to the untreated group. (D) Spot assays of *Candida albicans* grown at 37°C in the presence of chemical stressors alone or the mixture of chemical stressors and HCOS. Planktonic *C. albicans* cells in log phase were inoculated on the plate with ten-fold serial dilutions (starting with 1 × 10^5^ cells in the left column of each panel), treating with HCOS (125 μg/mL) alone or CFW (200 μg/mL), SDS (0.005%), NaCl (1 M) with or without co-incubation with HCOS (125 μg/mL). (E) SEM images of *C. albicans* cell treated with blank medium or 1mg/mL HCOS for 24 h. Scale bar, 3 μm. (F) TEM micrographs of *C. albicans* treated with blank medium or 1 mg/mL HCOS for 24 h. Scale bar, 200nm. ** p < 0.01, *** p < 0.001 and all experiments were repeated three times.

### The long-chain chitosan oligosaccharide (HCOS) was incorporated into the dynamic cell wall

To determine the binding site of HCOS on fungal cells, HCOS was labeled with the fluorescein FITC. The HCOS-FITC showed an unaffected antifungal activity as unlabeled HCOS (supplementary Fig. 1). Fluorescence confocal microscopic studies suggested that when fungal cells were exposed to a sub-IC50 concentration of HCOS-FITC, the HCOS-FITC was localized at the cell surface, colocalized with cell wall stains calcofluor white (CFW) and concanavalin A (ConA), but not inside the cell (Fig. 2A and B). On the contrary, LCOS mainly accumulated into the cell (Fig. 2A and B) and did not exhibit an apparent antifungal effect (Fig. 1A and B). Interestingly, time-dependent observations showed a fast and apparent accumulation of HCOS-FITC at apexes of the yeast cells after 5-minute exposure (Fig. 2C). HCOS stayed at the tip of the cell for at least 10 min and then distributed around the yeast cell surface (Fig. 2C). Enriched fluorescent signal in the chitin enriched alkali-insoluble cell wall extracts from HCOS-FITC treated cell further supposed that HCOS was integrated into the cell wall polysaccharide network (Fig. 2D). Besides, HCOS-FITC did not target the cell wall of the heat-killed *C. albicans* (supplementary Fig. 2). These results demonstrated that HCOS had an organized distribution pattern on the fungal cell wall.

**FIG 2.**
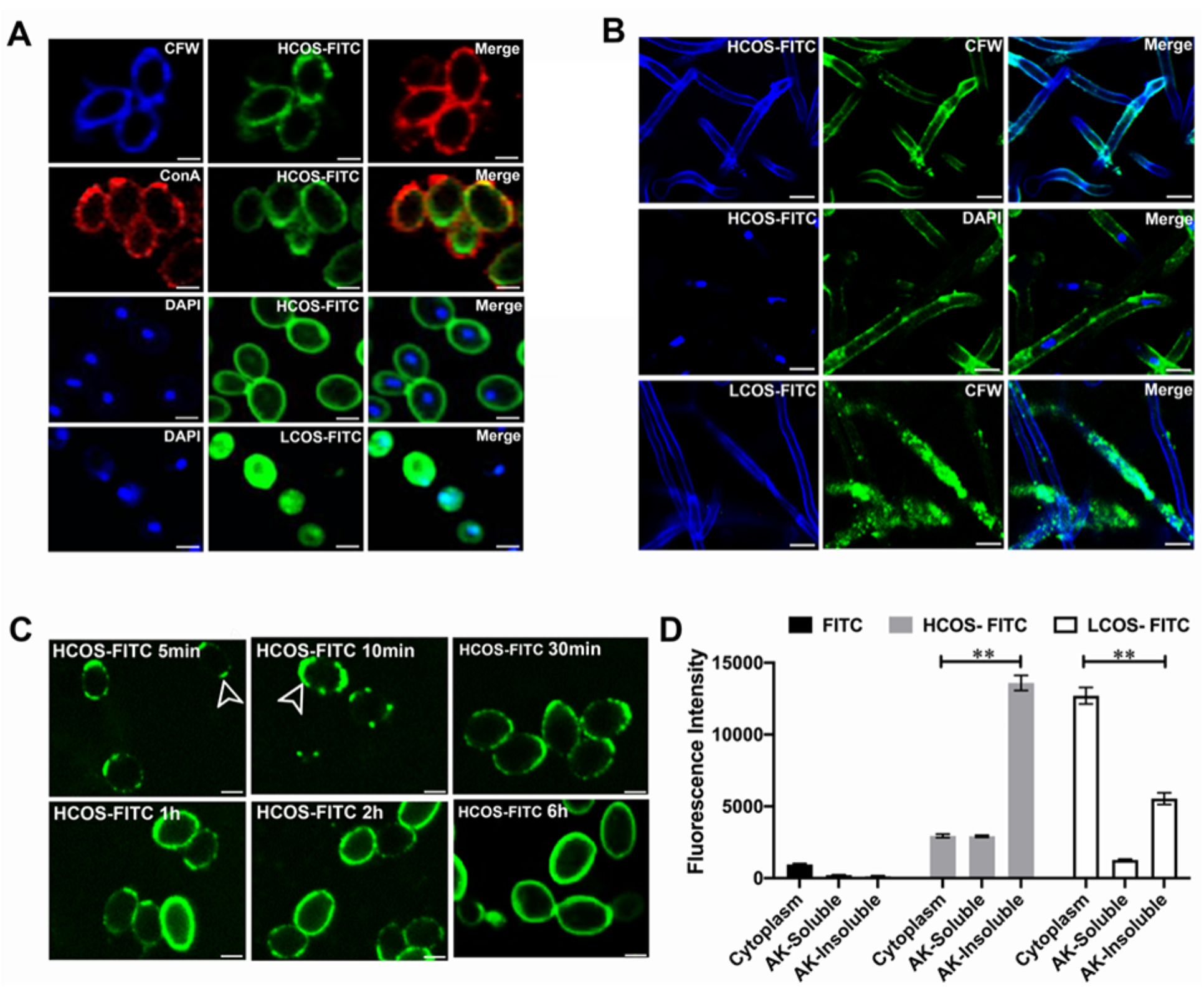
HCOS-FITC was incorporated into the dynamic cell wall. (A) Planktonic *C. albicans* were incubated with 500 μg/mL HCOS-FITC or HCOS-FITC for 24 h at 30°C. Then the cells were stained with 5 μg/mL CFW, 1μg/mL DAPI, or 10 μg/mL ConA. 5 μL stained cell samples were observed by fluorescence microscopy. Scale bar, 10 μm. (B) Biofilm exposure to HCOS-FITC (500 μg/mL) or LCOS-FITC (500 μg/mL) were stained with 5 μg/mL CFW and 1 μg/mL DAPI and observed under confocal microscopy. Scale bars, 10 μm. (C) Planktonic *C. albicans* were incubated with 500 μg/mL HCOS-FITC for 5 min - 6 h at 30°C, 5 μL samples were transferred onto a slide and observed under fluorescence microscopy. scale bar, 10 μm. (D) The cell wall of planktonic *C. albicans* was extracted with exposed with 500 μg/mL FITC, HCOS-FITC or LCOS-FITC for 6h, respectively. The fluorescence intensity of cytoplasm, alkali-soluble (AK-soluble) and alkali-insoluble (AK-insoluble) were measure. Data are represented as means ± SD (n=6). ** p < 0.01and all experiments were repeated three times.

### PHR family proteins are candidate targets of the long-chain chitosan oligosaccharide (HCOS)

To identify potential HCOS-binding targets, we set up an HCOS-affinity chromatography and screened proteins from fungal cell wall extracts (Fig. 3A and supplementary Fig.3A). Two protein bands around 22 kDa and 50 kDa was identified and processed for the LC-MS/MS-based protein identification (supplementary Fig. 3B). Among 146 candidates pulled out from MS/MS analysis (Supplementary Table. 1). Three candidates, PHR1, PHR2, and PGA4, drew significant attention. They all belong to the PHR β-(1,3)-glucanosyltransferases multigene family with transglucosidase activity, functioning in cell wall biosynthesis and morphogenesis. Besides, a parallel antifungal studies of combination usage of HCOS with different glucanases unexpectedly found that a *Fibrobacter Succinogens* β-(1,3)-glucanase impaired the antifungal activity of HCOS (supplementary Fig. 2C). Therefore, we primarily assess the potential relationship of PHR family proteins with HCOS. The binding between heterogeneously expressed Phr1p and HCOS was determined by microscale thermophoresis (MST) tests (Fig. 3B), showing a high affinity between Phr1p and HCOS with an equilibrium dissociation constant (KD) 3.22 μM (Fig. 3B) close to the KD of Phr1p and β-(1,3)-glucan (KD = 0.48 μM) (Fig. 3E). The KD of HCOS and Phr2p was 12.7 μM (Fig. 3C). Interestingly, the binding between HCOS and Pga4p was relatively weak (KD = 120 μM) (Fig. 3D) and the short-chain LCOS barely bound to Phr1p (KD = 3.7M) (Fig. 3F). To be noted, the RNA expression level of the *Phr1* gene was upregulated through the biofilm formation (Fig. 3G), consistent with the increased activity of HCOS against mature *C. albicans* biofilm (Fig. 1C).

**FIG 3.**
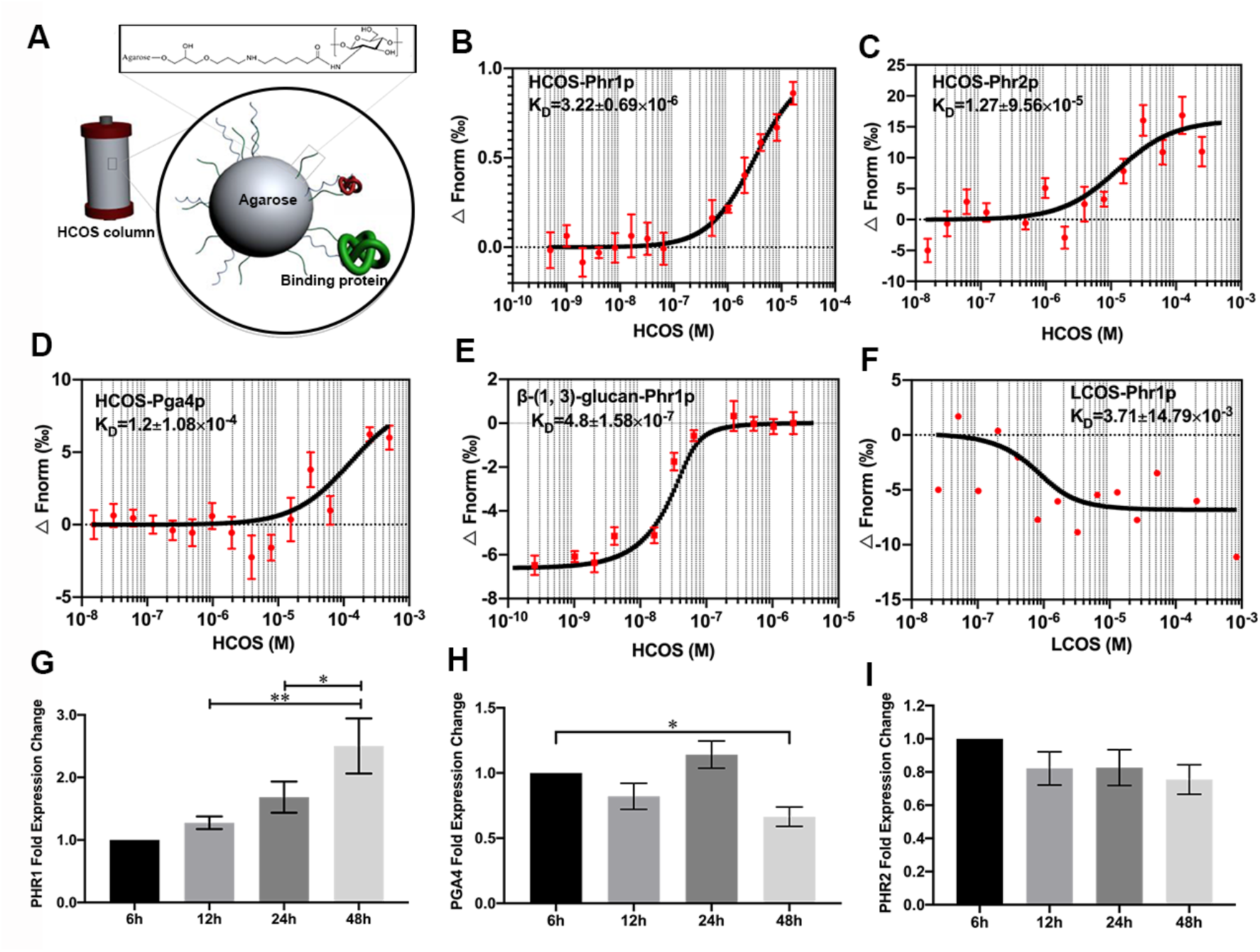
Identification of HCOS-binding proteins in the *C. albicans* cell wall. (A) HCOS was covalently bound to the NHS activated agarose via amine coupling to set up the HCOS column filler. Cell wall protein extract were loaded on the column to identify the HCOS-binding candidates, illustrated by green and red color molecules in the cartoon. (B) The MST curve showing the interaction of HCOS with Phr1p. (C) HCOS with Phr2p. (D) HCOS with Pga4p. (E) β-(1,3)-glucan with Phr1p. (F) LCOS with Phr1p was determined by employing standard data analysis with MO. Affinity Analysis Software. The graphs displayed data from three independent measurements. Error bars represented the SEM. (G) and (H) and (I)Relative fold change in the expression levels of *Phr1*, *Pga4* and *Phr2* in the different stages of biofilm formation was assessed by RT-PCR using *Act1* as reference. Data represented as means ± SD (n=4). * p < 0.05, ** p < 0.01 and all experiments were repeated three times.

### HCOS exerted its activity by interfering the β-(1,3)-glucanosyl transferases activity of Phr1p

To assess the influence of HCOS on the transglucosidase activity of Phr1p, laminarin was used as the donor substrate and an SR-labeled laminaripentaose (L5-SR) as the acceptor molecule in an enzyme assay *in vitro*. The transglycosylation efficiency of Phr1p was significantly inhibited by HCOS but not LCOS (Fig. 4A). However, HCOS could not act as a substrate for Phr1p (Fig. 4A). It suggested that HCOS acts as an inhibitor rather than a substrate for PHR transglucosidases. Consistently, a *Candida albicans Phr1* null mutant showed less sensitive to HCOS than the wild type (Fig. 4B). Fluorescent microscopic study also indicated that Phr1p was co-localized with HCOS-FITC (Fig. 4C). In the absence of Phr1p, HCOS-FITC still targeted the cell wall but did not showed the accumulation at cell apexes (Fig.4D). These results suggested that Phr1p was the main target of HCOS, while it is not fully responsible for the integration of HCOS into the cell wall.

**FIG 4.**
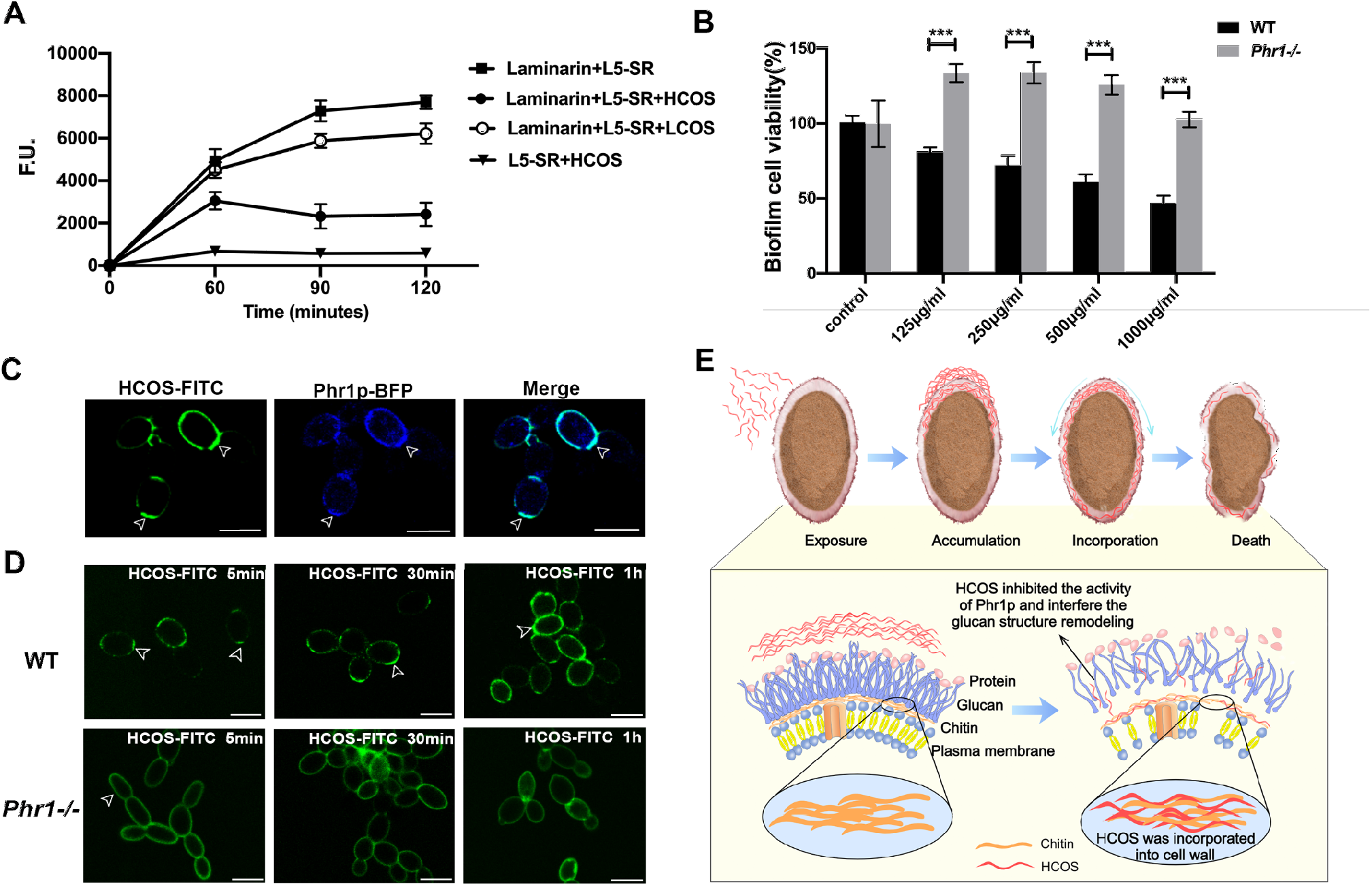
HCOS interfered the β-(1,3)-glucanosyl transferases activity of Phr1p. (A) Phr1p transglycosylation activity were deterimined using equimolar concentrations of the SR-labelled laminaripentaose and laminarin as substrate, with or without 1 mg/mL HCOS or LCOS. Data represented as means ± SD (n=3). (B) Susceptibility of a wild type *C. albicans* strain (GH1013) and a PHR1 null mutant *Phr1−/− t*o HCOS. The biofilm cell viability was determined by XTT assay and normalized to the control. Data represented as means ± SD (n=6). (C) Confocal micrographs of *C. albicans* strain expressing Phr1p-BFP (blue) treated with 500 μg/mL HCOS-FITC for 2 h. (D) Planktonic wild type *C. albicans* and *Phr1* null mutant were incubated with 500 μg/mL HCOS-FITC for 5 min, 30min and 1h at 30°C, 5 μL samples were transferred onto a slide and observed under fluorescence microscopy. scale bar, 10 μm. (E) Schematic illustration of HCOS antifungal action. ** p < 0.01 and all experiments were repeated three times.

### Synergistic effects of HCOS with antifungal drugs against *Candida* species

HCOS greatly affected the integrity of the cell wall, which provides a critical protective barrier to the fungi, and HCOS acted well on established biofilm (Fig.1). These findings indicated its great potential as an antifungal drug synergist, especially against fungal biofilm. We assessed the synergetic activity of HCOS with different antifungal drugs against *C. albicans* biofilm. After exposure to 500 μg/mL HCOS together with 1000 μg/mL fluconazole or 1 μg/mL caspofungin, the fungal biofilm mass was decreased by 57% (FLC+HCOS) or 44% (caspofungin+HCOS), respectively (Fig. 5A and C). Moreover, the microbial viability was decreased dramatically by 93% (FLC+HCOS) or 80% (caspofungin+HCOS) (Fig. 5B and D). Either drug alone showed no obvious impact on the fungal biofilm at the same concentration. Combining HCOS and antifungal drugs showed a strong anti-biofilm effect after 1 h treatment and efficiently eliminated biofilm cells with prolonged treatment time (Fig. 5E and F), resulting in the loose and rough cell surface, shrunk and ruptured cell shape (Fig. 5H). The results were further confirmed by a fluorescence-based live-dead assay and a microfluidic biofilm model (supplementary Fig. 6A). We then systematically evaluated the synergistic activity of HCOS with drugs against planktonic or biofilm cells of *C. albicans* and various other *Candida* strains, including *Candida tropicalis*, *Candida parapsilosis*, *Candida glabrata*, and *Candida rugosa*, using the standard checkerboard titration method. The result showed a significant synergistic effect between the drugs and HCOS against all tested strains based on the fractional inhibitory concentration index (FICI) model, as the FICI values in the range of 0.063 to 0.5 (Table 1 and supplementary Table 2).

**Table 1.**
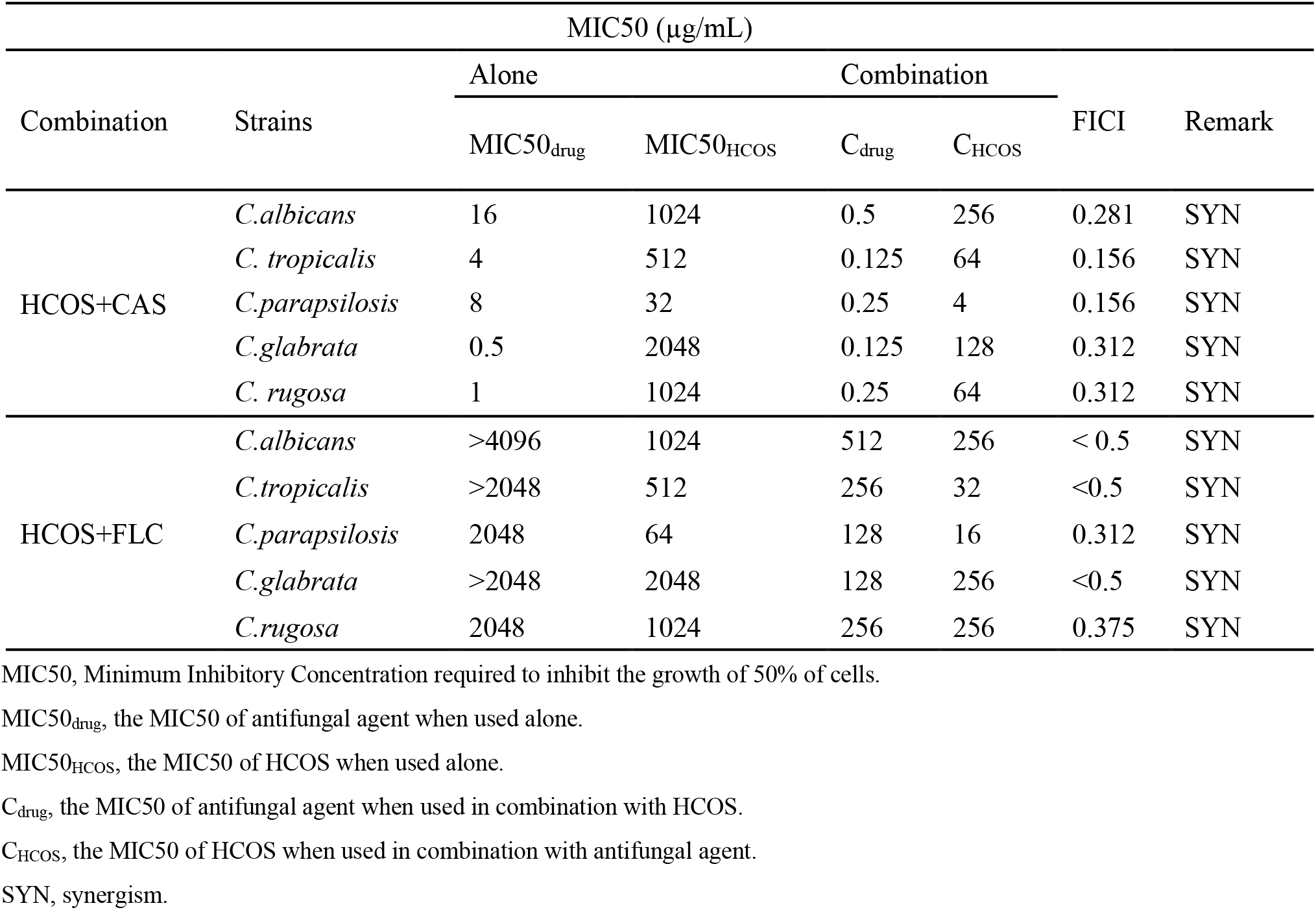
Fractional inhibitory concentration index (FICI) of combination between antifungal drugs and HCOS against *Candida* biofilms.

**FIG 5.**
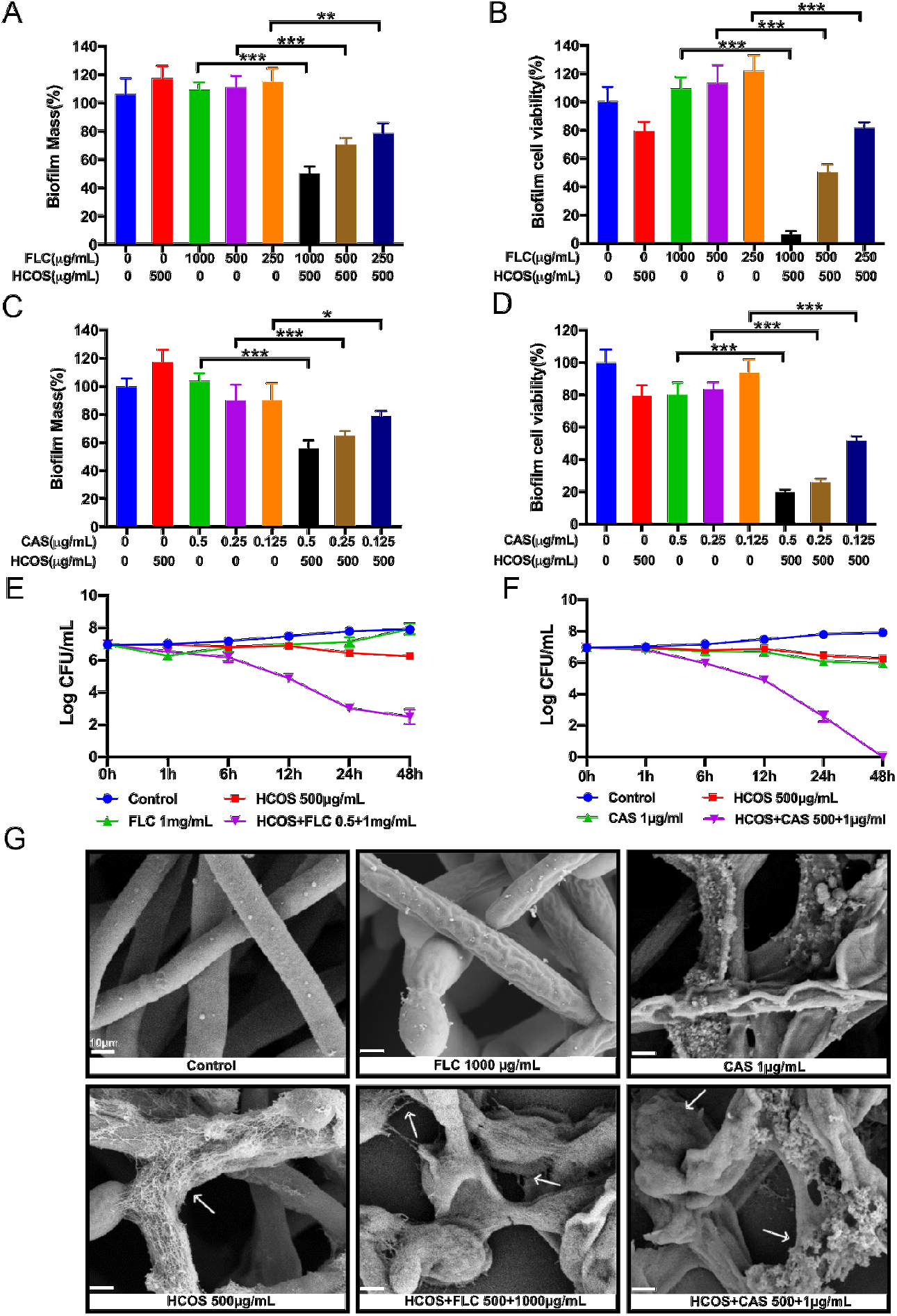
The synergistic antifungal effects of HCOS combined with the antifungal drug against *Candida albicans* biofilm. (A) and (C) The effects on *C. albicans* biofilm mass of fluconazole and caspofugin alone or in combination with 500 μg/mL HCOS. Y-axis represents the value of the ratio of biofilm mass in treated samples relative to the control. (B) and (D) The effects on the biofilm cell viability of fluconazole and caspofugin alone and in combination with 500 μg/mL HCOS. Y-axis represents the value of the ratio of viable cells in treated samples relative to the control. (E) and (F) Mature *C. albicans* biofilms were exposed to HCOS (500 μg/mL), fluconazole (1 mg/mL), caspofugin (1 μg/mL) or the combination of the drug and HCOS for 1 h, 6 h, 12 h, 24 h and 48 h. Biofilm incubated with plank medium was used as a control. Aliquots were obtained at the indicated time points and serially dilutions were spread on agar plates. Colony counts were determined after 48 h of incubation. The biofilm mass was determined using crystal violet assay and biofilm cell viability was determined using XTT reduction assay. Data are shown as the means ± standard deviations for three independent experiments. ∗ P < 0.05, ∗∗∗ P < 0.001 represent the difference between the indicated group. (G) Mature biofilm exposured to 1000 μg/mL FLC,1 μg/mL CAS, 500 μg/mL HCOS, HCOS + FLC (500+1000 μg/mL), HCOS +CAS (500+1 μg/mL) for 24 h and observed under scanning electron microscopy.

### The antifungal activity of the combination of HCOS and fungicide in a murine model of systemic candidiasis

To further evaluate the efficacy of combination therapy with fungicide and HCOS, we compared the survival rates and renal fungal burden and renal tissue sections of *C. albicans*-infected mice using the murine model of systemic candidiasis. Firstly, we checked the *in vitro* human cells toxicity of HCOS and found that HCOS had neither significant toxic effect on HUVECs nor RAW 264.7 at concentrations that show significant antifungal activity (Fig.6A and B). In the murine model of systemic candidiasis, treatment of mice with combination of 50 mg/kg HCOS and 0.1 mg/kg fluconazole showed significantly strikingly increased survival rates (Fig. 6C). All the mice survived after treatment with HCOS (50 mg/kg) combined with fluconazole (0.1 mg/kg) within 7 days, superior to fluconazole alone (0.1 mg/kg, survival rate of 20%) or HCOS alone (50 mg/kg, survival rate of 40%). Consistently, the renal fungal burden of *C. albicans* infected mice significantly reduced under combinatorial therapies within 7 days while fluconazole or HCOS alone only showed a mild effect (Fig. 6D). PAS-stained histological examinations of kidneys also showed that the combined use of HCOS and fluconazole markedly reduced the number of fungal cells, especially hyphae and pseudohyphae (Fig. 6E).

**FIG 6.**
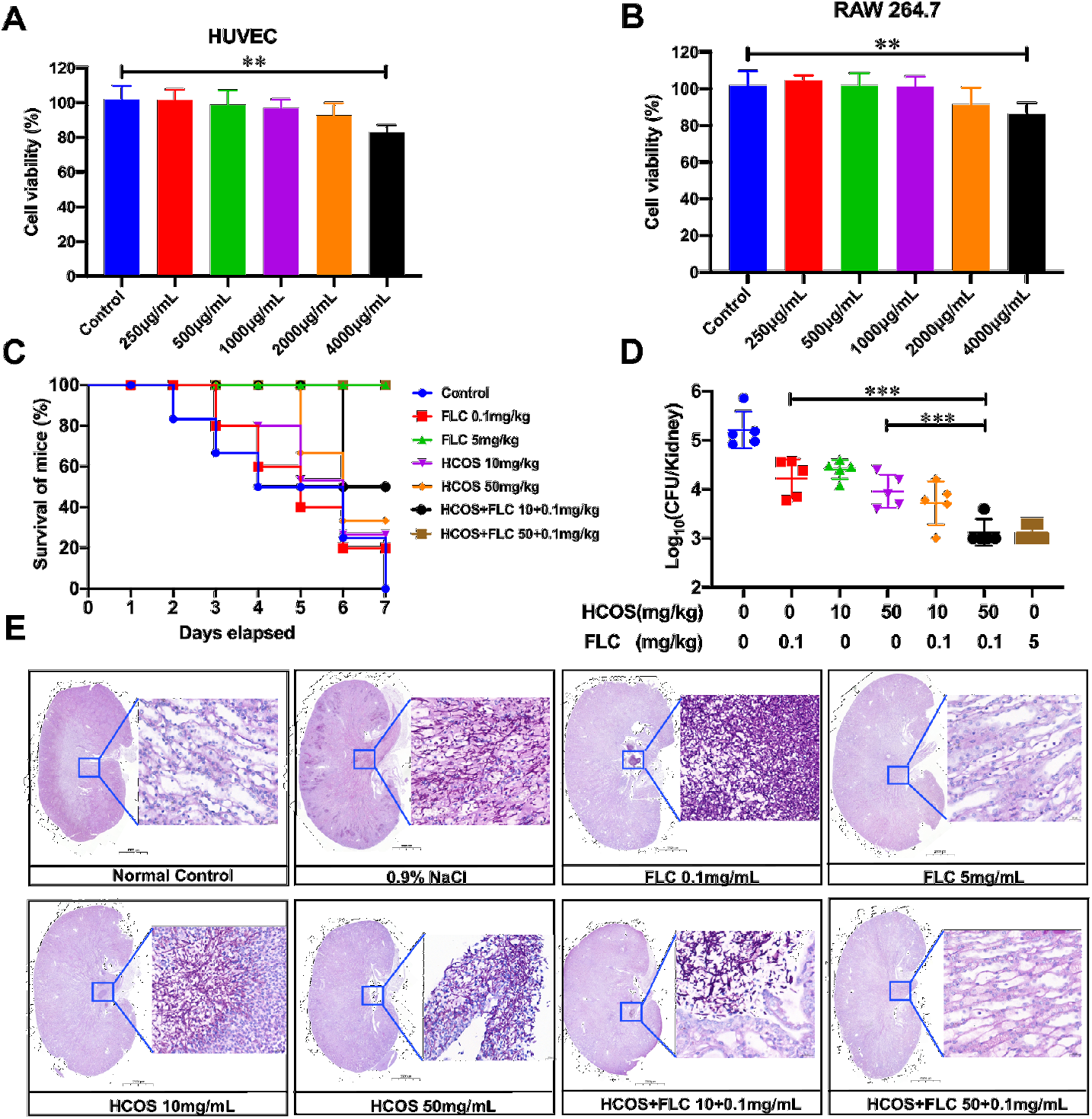
The combination treatment of HCOS and FLC on the murine model of systemic candidiasis. The cytotoxic effect of HCOS on HUVEC (A) and RAW 264.7 (B) viability was assessed by the MTT test following 24 h treatment. (C) Survival rate of infected mice (n = 5) treated with fluconazole at the concentration of 0.1 mg/mL or 5 mg/mL alone by intraperitoneal injection (ip), HCOS at the concentration of 10mg/mL or 50 mg/mL alone by intragastrical administration (ig), or treated with the combination of HCOS and fluconazole at levels of 10+0.1 mg/mL and 50+0.1 mg/mL. Survival rate was observed after 7 days treatment. The log-rank test was used for statistical analysis. (D) The fungal burden of the kidney after antifungal treatment (in log_10_ CFU/mL of kidney) was plotted. Data are shown as the means ± SD (n=5). (E) PAS staining of the kidneys. *Candida* hyphae were stained with purple red color. The scale bar of kidney is 2mm, the scale bar of magnified images of areas inside the blue boxes is 20 μm. ∗∗P< 0.01, ∗∗∗ P < 0.001 experiments were repeated three times.

## DISCUSSION

The fungal cell wall is a protective barrier that is critical for cell viability, making it an attractive target for the development of antifungal therapies, especially because mammalian cells have no equivalent structures (6, 27). Compounds inhibiting cell wall polysaccharide biosynthesis, such as echinocandins and nikkomycins, have been developed, and some of them became crucial first-line drugs for antifungal treatment in clinical practice (28, 29). In this study, a highly deacetylated long-chain chitosan oligosaccharide HCOS mimicking cell wall chitin/chitosan fragments were used to test whether it can sneak into the cell wall remodeling process and interfere its integrity. Results showed that HCOS was promptly incorporated into the *Candida* cell wall, disturbed the wall morphology, and presented apparent antifungal activity on both planktons and biofilm (Fig. 1 and Fig. 2). HCOS exerted its activity most probably by interfering with family members of PHR β-(1,3)-glucanosyl transferases (Fig. 4). Furthermore, HCOS showed great synergistic activity with different fungicides against both *Candida* planktons and biofilm, especially those in biofilm (Fig. 5 and Fig. 6, Table1 and Supplementary Table 2), likely due to its effect on the cell wall integrity and unusual enhanced activity against established biofilm. These findings indicated HCOS has a great potential as an antifungal drug synergist, especially against fungal biofilm.

Apexes are the most active sites of cell wall remodeling and chitin synthesis (8, 30), where enzymes responsible for the cell wall biosynthesis were concentrated (31). When treated to the *Candida* cell, the HCOS was fast accumulated at apexes of the yeast cells and septa of the pseudohyphae, and then distributed around the cell surface (Fig. 2C). However, HCOS-FITC was not accumulated at the cell wall of the heat-killed *C. albicans* (supplementary Fig. 2). It suggested that cell wall proteins were critical for the binding of HCOS to the cell wall. Also, HCOS was mainly in the alkali-insoluble chitin potion of cell wall exacts (Fig. 2D). The alkali-insoluble portion from HCOS-treated cell wall exacts showed an increased ratio of glucosamine compared to untreated cells (supplementary Fig. 4). These observations suggested that HCOS sneaked into the cell wall chitin/chitosan organization process. The oligosaccharide size is critical for this action because of a short chain chitosan oligosaccharide LCOS accumulated into the cell instead of the cell surface (Fig. 2A and B). Also, LCOS did not exert noticeable antifungal effects (Fig. 1A and B). Previous studies suggested that the chitosan polysaccharide exerted antifungal activity by targeting plasma membrane (32–35). Therefore, incorporation with the cell wall was the key to the antifungal activity of the chitosan oligosaccharide, which highly depends on the oligomer’s size, not too short, not too long.

Using affinity chromatography, we tried to isolate and identify *C. albicans* cell wall proteins binding HCOS. We initially use Zymolyase-treated cell samples to isolate HCOS-binding proteins, but did not find any proteins bound to the affinity column (data not shown). We then added the GPI phospholipase together with Zymolyase to treat cells. Multiple bands were eluted from the column (supplementary Fig. 3A). Two protein bands around 22 kDa and 50 kDa, which was enriched in the elution but not the flow-through, was processed for the LC-MS/MS-based protein identification in the first place. our microscopic studies showed that HCOS always stays on the cell surface. Therefore, HCOS’s targets likely are GPI-anchored proteins and present on the cell surface. There are three homogenous proteins, PHR1, PHR2, and PGA4, among 146 candidates from peptide mass fingerprinting (Fig. 3A and Supplementary Table 1) deserved more attention. These three proteins are located on the cell wall and have a molecular weight around 50 kDa after removal of the GPI anchor. They belong to the PHR multigene family of β-(1, 3)-glucanosyltransferases and have been classified in the GH72 family in the Carbohydrate Active Enzyme database (http://www.cazy.org/) (36). Interestingly, a parallel antifungal studies showed that a *Fibrobacter Succinogens* β-(1,3)-glucanase impaired the antifungal activity of HCOS (see Fig. S3C in the supplementary material), indicating the potential relationship with β-(1,3)-glucanase and HCOS. Our experiments showed that HCOS had a high affinity to Phr1p. The C-terminal domain of Phr1p might be necessary for interaction with HCOS, considering the binding between HCOS and Pga4p was relatively weak (Fig. 3D). *In vitro* enzymatic analysis showed that HCOS inhibited the transglucosylase activity of Phr1p (Fig. 4A). Also, The *Phr1* but not *Phr2* null mutant showed resistant to HCOS (Fig. 4B and see Fig. S5 in the supplementary material). These results suggested that HCOS might interfere with the cell wall glucan modeling through Phr1 proteins (Fig. 4E). We noticed that many intracellular mitochondria and nuclear proteins were present in the table of mass-based candidate binding proteins of HCOS. It is possible that chitosan oligosaccharides could bind intracellular proteins, if they could get into the cell, such as LCOS. However, since our results showed that HCOS always stay at the cell surface, these intracellular proteins are unlikely involved in the antifungal activity of HCOS. The presence of intracellular proteins in the HCOS-binding list might be due to the application of GPI-specific phospholipase and Zymolyase, which might increase the chance of cell-leaking during the experiment.

PHR proteins act as cell wall remodeling enzymes and play an active role in cell wall biosynthesis (36), glucan elongation and branching (37). Enzymatic analysis showed that HCOS acts as an inhibitor rather than a substrate for Phr1p transglucosidases (Fig. 4A). Fluorescence microscope results showed that HCOS could still bind to the cell wall of *Phr1* null mutant (Fig. 4D). However, HCOS was not enriched at the tip of the *Phr1* null cell, unlike its localization on the wild type of cell surface. We also hydrolyzed the cell wall of HCOS treated cells and analyzed the component of degraded cell walls, including chitin/chitosan fragments, glucan fragments, and chitin-glucan fragments. We did not see obvious difference in the composition of the degradation products between HCOS treated and untreated samples (data not shown). Therefore, these results suggested that HCOS accumulated in the cell wall via non-covalent interactions. PHR1 was important for the apex distribution and antifungal activity of HCOS but dispensable for the cell wall binding of HCOS. It is possible that other proteins might be involved in the action besides PHR proteins. Other HCOS-binding candidates and enzymes involved in chitin-glucan and chitin-chitin cell wall crosslinks, such as CRH family proteins (38–40), were worth for further investigations. On the other hand, we could not rule out the possibility that HCOS accumulate in the cell wall via covalent interactions.

A possible explanation for the action of HCOS on cell wall organizing machinery is that HCOS is structurally similar but not identical to the cell wall chitin/chitosan fragments. HCOS might be recognized as nascent chitin/chitosan polymers by cell wall remodeling enzymes. Although chitin was partially deacetylated in the cell wall, study showed a non‐uniform deacetylation pattern of chitin in fungal cell wall (41). Instead, HCOS had a continuously super deacetylation pattern. The structural difference caused by deacetylation might help HCOS interfere the activity of chitin/chitosan related enzymes. Here, our findings support an unexpected tight relation between chitin deacetylation and glucan branching and elongation, although the detail of chitin deacetylation on the cell wall remains unclear.

The RNA expression level of the PHR1 gene was upregulated through the maturation of the *Candida* biofilm (Fig. 3G-I). It explains the enhanced activity of HCOS against the mature biofilm (Fig. 1C). Usually, the maturation of biofilm increases the antifungal resistance of fungal cells (42). Surprisingly, the fungicidal activity of HCOS became more robust following the maturation of the biofilm (Fig. 1C). Moreover, a combination of first-line antifungal drugs and HCOS effectively killed planktons and eliminated fungal biofilms of several common pathogenic *Candida* species (Table 1 and Supplementary Table 2), suggesting a synergistic activity of HCOS and fungicides. Antifungal treatment on biofilm-associated fungal infections is often ineffective, leading to recurrent and chronic infections and biofilm-specific drugs are not available currently (42). A combination of HCOS and fluconazole significantly increased the survival rate and reduced the renal fungal burden of *C. albicans* infected mice (Fig. 6). These results strongly suggested that HCOS had excellent potential for developing biofilm-specific antifungal drugs or drug synergists. Although it required a relative high concentration of HCOS to exhibit antifungal activity, high concentration HCOS did not show obvious cell toxicity to mammalian cells (Fig.6A and B) (43). Several studies showed that as high as 2000 mg/kg/day orally administered chitooligosaccharide showed no toxicity in the rat (44). Thus, chitooligosaccharide is much safer than commonly used antifungal drugs. To be noted, our animal studies showed that 50mg/kg HCOS alone already showed noticeable improvement in the survival rate. Natural glycan-based drugs might need relatively high concentration to treat the pathogenic infection. However, they usually do not have high toxicity and strong side effect. Also, HCOS showed a significant synergist effect with other fungicides (Table 1, Table S2). In this case, the effective concentration of both HCOS (around 100 μg/ml) and the drug is lower. In the antibiofilm assay, HCOS alone even showed a lower MIC value compared to fluconazole. In fact, chitosan and chitosan oligosaccharide have been widely used as foods and biomedical materials (45, 46). It has been proved with good biocompatibility and safety to human body. Moreover, HCOS might be used as a carrier of other cell wall acting compounds or further modified to improve the antifungal activity. This study opens new therapeutic perspectives for treating human candidiasis.

## METHODS

### Materials and strains

The high molecular weight chitosan oligosaccharide lactate (HCOS) (MW= 4,000-6,000, average MW 5,000) was purchased from Sigma (St. Louis, MO, USA). The low molecular weight chitosan oligosaccharide lactate (LCOS) (average MW: 835 Da) was purchased from GlycoBio (GlycoBio, Dalian, China). Fluconazole (FLC) and caspofungin (CAS) were purchased from Sigma (St. Louis, MO, USA). Unless specified, all other chemicals or reagents were obtained from Sigma and Solarbio (Beijing, China).

HCOS-FITC conjugate was prepared as previously described with minor modifications (47, 48). Chitosan oligosaccharides HCOS or LCOS (0.2 g) were dissolved in 20 mL of H2O and adjusted to pH 6.8 with NaOH. Fluorescein Isothiocyanate (FITC) was then added and reacted at room temperature under stirring for 5 h. COS-FITC conjugates were precipitated with ethanol and rinsed with 70 % methanol repeatedly to remove unattached FITC. The precipitate was collected after lyophilization.

*Candida albicans* (SC5314), *Candida parapsilosis* (ATCC22019), *Candida tropicalis* (ATCC7349), *Candida glabrata* (ATCC2001), *Candida rugosa* (ATCC2142) were purchased from China General Microbiological Culture Collection Center (CGMCC). **Δ*Phr*1/Δ*Phr*1** strain and **Δ*Phr*2/Δ*Phr*2** used in the experiment were generously granted by Prof. Huang guanghua.

The plasmid backbone for PHR1-GFP expressing was derived from pENO1-BFP-NAT (Addgene), the BFP sequence was inserted between the amino acids G489 and G490 of PHR1(49). The recombinant plasmid was transformed into *C. albicans* by electroporation, and the successfully transformed clone was screened by 400 μg/mL nourseothricin (NAT) (Solarbio, Beijing, China).

All *Candida* strains were grown at 30 °C if not otherwise indicated. Cell were grown on yeast extract-peptone-dextrose (YPD) agar or cultured in YPD medium (1% yeast extract, 2% peptone, and 2% dextrose) with shaking (200 rpm) for 14-16 h. For biofilm seeding, overnight cultured cells were adjusted to the desired cell density and cultured in RPMI-1640 medium supplemented with L-Glutamine, without sodium bicarbonate (Thermo Fisher, Waltham, USA) and buffered with morpholine propanesulfon (MOPS) to pH 7.0.

### Mice

6-8 weeks old Kunming female mice were obtained from Beijing HuaFuKang Biotechnology company. All mice weighed 25-30 g when tested. Mice were adapted to standardized environmental conditions (temperature = 23 ± 2 °C; humidity = 55 ± 10%) for 1 week before infection. Mice were maintained in strict accordance with the regulations for the Administration of Affairs Concerning Experimental Animals approved by the State Council of People’s Republic of China (14 November 1988). The laboratory animal usage license number is SCXK-2019-0008, certified by Beijing Association for Science and Technology.

### Antifungal susceptibility testing of *Candida* cells

The antifungal activity of HCOS, fungicides, and their mixture was determined by a broth microdilution method in 96-well culture plates comply with the Clinical and Laboratory Standards Institute (CLSI) guidelines outlined in document M27-A3 (50). The MIC50 was defined as the lowest drug concentration corresponding to a 50% reduction in cell growth. All experiments were performed in triplicate. The synergistic interaction of pairwise drugs was determined based on the fractional inhibitory concentration index (FICI) model (51). FICI values of ≤ 0.5, 0.5 < FICI ≤ 4.0 and > 4.0 represent synergism, no interaction and antagonism, respectively.

To assess antifungal susceptibility of *Candida* biofilm, *Candida* biofilms were established in 96-well polystyrene microtiter plates as previously described (52, 53). Briefly, 100 μL of *Candida* cells (1 × 10^7^ cells/mL) in RPMI-1640 medium were added to a 96-well polystyrene microtiter plate and cultured for 24 h at 37 ℃ to allow biofilm formation. Mature biofilms were washed with PBS and replenished with 100 μL fresh RPMI-1640 medium containing different drug concentrations. The plates were then incubated at 37℃ for a further 24 h. The biofilm mass was determined by crystal violet assay (54). The sample was measured for absorbance at 590 nm with a TECAN Infinite M200 PRO multifunction microplate reader (TECAN, Groding, Austria).

Fungal cell viability was quantitatively assessed using a 2,3-bis-(2-methoxy-4-nitro-5-sulfophenyl)-2H-tetrazolium-5-carboxanilide (XTT) reduction assay, as previously described (55). Briefly, 90 μL of XTT (1 mg/mL) and 10 μL phenazine methosulfate (320 μg/mL) were added to each well. The plate was then incubated at 37 °C for 2 h. The sample was measured for absorbance at 490 nm using a TECAN Infinite M200 PRO multifunction microplate reader (TECAN, Grodig, Austria). All tests were performed in six replicates for each treatment. Each assay was performed with three biological repeats.

### Fluorescence confocal laser scanning microscopy

*C. albicans* cells from an overnight culture were adjusted to 1×10^7^ cells/mL with fresh YPD medium. HCOS-FITC or FITC with the concentration of 500 μg/mL were added into the sample and incubated for 5 min,10 min, 30 min, 1 h, 2 h, and 24 h, respectively. Then washed cell samples were spotted in 5 μL aliquots on a glass microscope slide for the fluorescence microscopy study. Heat-killed *C. albicans* cells were prepared by incubation at 121°C for 30 min. Mature *C. albicans* biofilm cultured on sterile coverslips were treated with indicated drugs at 37 ℃ for 24 h. The biofilms were then washed and fixed using a 4% paraformaldehyde solution for 30 min at 30 °C. Fixed samples were stained with 1μg/mL DAPI, 10 μg/mL ConA, 5μM CFW or 1μg/mL PI for 30 min at 30 °C. After washing, the coverslip was placed and sealed on a slide. Stained samples were observed under LEICA CTR4000 confocal laser scanning microscope (Leica, Barnack, Germany).

### Scanning electron microscope (SEM)

SEM was performed as described in reference (56) with minor modifications. *C. albicans* cells were cultured on sterile coverslips in 24-well plates. After antifungal treatment, samples were washed with PBS and fixed in 4% paraformaldehyde with 2% glutaraldehyde in 0.1M phosphate buffer for 45 min at 4 ℃. Samples were then briefly rinsed in 0.1M phosphate buffer before post-fixing with 1% OsO4 (Sigma-Aldrich, St. Louis, MO, USA) for 45 min at ambient temperature. Then the samples were rinsed in 0.1M phosphate buffer and gradually dehydrated using a series of increasing concentrations of ethanol in water (50, 70, 80, 90, 95, 100%), followed with critical point drying. The coverslips were then mounted, and gold coated. Images were obtained with a scanning electron microscope (SU8010) at 5 kV.

### High-pressure freeze substitution transmission electron microscopy (HPF-TEM)

HPF-TEM was performed as described previously (57, 58). Living cell culture was collected by centrifugation in a microcentrifuge for 2min. The pellets were resuspended in 2% agarose and transferred to flat specimen carriers. Following HPF, the fast-frozen samples were immersed into a freezing tube containing 2% osmium tetroxide in 100% acetone and placed into the freeze-substitution device (Leica EM AFS, Germany) with the following parameters: T1 = −90°C for 68 h, S1 = 3°C/h, T2 =−0°C for 12 h, S2 = 3°C/h, T3 = −30°C for 10 h, then slowly warmed to 10°C (3°C/h). Following FS, four rinses in 100% acetone, 15 min each, at room temperature (rt), then transfer samples into a new 2ml Eppendorf tube. After that, Samples were infiltrated in graded mixtures (1:3, 1:1, 3:1) of resin (EMS, Resin Mixture: 19.6 ml SPI-PON812, 6.6 ml DDSA, 13.8 ml NMA,1.5%BDMA) and acetone mixture, then changed 100% resin 5 times for 3 days on a rotator. Finally, samples were embedded and polymerized 12h at 45°C and 48h at 60°C. The ultrathin sections (70nm thick) were sectioned with a microtome (Leica EM UC7), double-stained by uranyl acetate and lead citrate, and examined by a transmission electron microscope (FEI Tecnai Spirit120kV).

### HCOS affinity chromatography of *C. albicans* cell wall proteins

The HCOS affinity column was prepared as described previously (59). An aliquot of 100 mg HCOS was dissolved in 10 mL 0.1 M NaHCO3-Na2CO3 buffer (pH 8.3), then added into 10 mL NHS-activated agarose. After the reaction mixture was shaken (50 rpm) at 4 °C for 14 h, the product was washed with 20% ethyl alcohol then packed into a 1.5 × 5 cm column.

Extraction of cell wall proteins was processed as previously described (60). Briefly, *C. albicans* biofilms were dislodged and gently sonicated at 42 kHz for 20 min with Branson 1510 Ultrasonic Cleaner sonicator to release biofilm cells without cell wall disruption followed by sonication at an amplitude of 70 for 10 min using an Intrasonic Processor (Cole Parmer, Vernon Hills, IL). The sample was then centrifuged and resuspended in 20 mM Tris-HCl (pH 7.0). Enzymolysis was performed using 0.1 mg/mL GPI specific phospholipase (generously granted by Prof. Guan Feng) and 50 U/g Zymolyase 20T at 30 °C for 3 h with gentle shaking. After centrifugation, the supernatant was collected and stored at 4 °C for further purification. The HCOS affinity column was pretreated with 1% OVA solution and successively washed with 20 mM Tris-HCl buffer (pH 7.0). After the sample was loaded, the column was equilibrated with 20 mM Tris-HCl buffer and eluted with 0.17 M Glycine-HCl buffer (pH 2.3). Eluted peak fraction was polled in tubes added with a 1/10 volume of 1 M Tris for immediate neutralization (61). The collected fractions were further separated by SDS-PAGE. The target band of the HCOS binding proteins was cut off from 12% SDS-PAGE gel and digested with trypsin. The digested sample was analyzed by LC-MS/MS (Orbitrap Fusion, Thermo Fisher scientific, USA). The Mascot daemon and Mascot server (V 2.5, Matrix science, Boston, USA) were used for the acquisition of MS data. The MS/MS data was analyzed with Proteome Discoverer (version1.4.0.288, Thermo Fischer Scientific) using Uniprot proteome-*Candida albicans* 2019 database.

### Cloning, expression and purification of Phr1

N-terminal GST tagged recombinant proteins containing a truncated version of the *C. albicans Phr1* and *Pga4* gene without the sequence encoding signal peptide and the GPI anchor were constructed using primers listed in Supplementary Table 3 and cloned into the bacterial expression vector pGEX6P1 (GE healthcare, Stockholm, Sweden). Plasmids were then transformed into *E. coli BL21* (DE3) cells grown in LB medium with 100 μg/mL ampicillin at 37°C until OD600 reached 0.6-0.8. IPTG with a concentration of 0.1 mM was added, and the culture was incubated at 16 °C for 24 h. The protein was then purified by GST Sefinose™ Resin kit (Sangon Biotech, Shanghai, China).

### Microscale thermophoresis (MST) assessment

Phr1, Phr2 and Pga4 proteins and Bull Serum Albumin (BSA) were labeled using the Monolith™ RED-NHS Protein Labeling Kit (NanoTemper Technologies) according to the manufacturer’s instructions. 10 μM labeled proteins were used as binding targets, while HCOS, LCOS or β-(1,3)-glucan (Putus Macromolecular, Wuhan, China) as binding ligands was titrated in 1:1 dilution series (concentrations starting from 1000 μM to 15 nM). Samples were loaded in the Monolith NT.115 MST standard-treated capillaries (NanoTemper Technologies), mixed well quickly, and immediately measured by MST (NanoTemper Technologies) at 37°C. All the experimental parameters used by the MST instrument were adjusted to 20% LED power and 40% MST power. Laser on and off times was set at 30 and 5 s, respectively. Triplicates of independently pipetted measurements were analyzed. The MO-Affinity Analysis software (version 2.1.3, Nano Temper Technologies) was used to determine the binding affinity by K_D_ value.

### Enzymatic assay

To determine the β-1,3-glucanosyltransferase activity of the Phr1 protein, we adopted the previously described fluorescent assay (62, 63). Fluorescent labeling of the laminaripentaose (Megazyme, Bray, Ireland) was accomplished by reacting the respective oligosaccharide glycamines with Lissamine rhodamine B sulfonyl chloride (LRSC) (Acros Organics, Belgium, USA) as described previously (64). The reaction mixture contained 1 mg/mL laminarin (Solarbio, Beijing, China), 30 μM SR-labeled laminaripentaose(L5-SR), 10 μg of Phr1p in 50 mM citrate-phosphate buffer at pH 5.5. 1 mg/mL HCOS or LCOS was added to the reaction to observe their effect on the enzymatic activity of Phr1p. Reactions were carried out at 30 °C for 0, 60, 90, and 120 min. The reactions were stopped by addition with 20 μl 40% (v/v) formic acid. 5 μl aliquots were spotted in triplicates on to Whatman 3 mm chromatographic paper and dried. The paper was then washed for 8-16 h with three changes of 5% formic acid in 66% (v/v) aqueous ethanol. The paper was dried, and photos were taken under the UV lamp. The fluorescent intensity of the spots was measured with Image J 1.53.

### Degree of deacetylation of the cell wall chitin by HPLC analysis

The cell wall of *C. albicans* was extracted as previously described (65). *C. albicans* were grown in YPD to mid-exponential phase and treated with HCOS, LCOS for 2 h at the concentration of 500 μg/mL, respectively. Cells were washed with sterile water and lysed with liquid nitrogen extraction thoroughly. The pellets were washed with 1 M NaCl repeatedly to remove contaminating cytoplasmic proteins and freeze-dried. The cell wall (20 mg) was hydrolyzed with 2 M trifluoroacetic acid for 3 h at 120 °C and freeze-dried after concentrating by rotary evaporation. The samples were resuspended in water to a 10 mg/ml concentration and analyzed by HPLC as described previously (66), and the relative proportions of monosaccharides in the cell wall were calculated.

### Distribution of COS in the cell wall

The cells were treated with HCOS-FITC and LCOS-FITC for 2 h at the concentration of 500 μg/mL to identify the cell wall components that HCOS incorporates. The cell wall extraction method is as described above. After freeze-dried, the samples were suspended in 1 M sodium hydroxide and heated again at 80 °C for 1 h and centrifuged. The fluorescent signals of the precipitate and supernatant were measured by TECAN Infinite M200 PRO multifunction microplate reader (TECAN, Grodig, Austria).

### Toxicity evaluation of HCOS

Human umbilical vein endothelial cells (HUVECs) and RAW 264.7 cells were obtained from the American Type Culture Collection (Manassas, VA, USA). Cells were grown in DMEM containing 10% FBS and 100 units/mL penicillin under 5% CO_2_ atmosphere at 37 °C. Cells were incubated in 96-well plates (3 × 10^3^ cells/well) with HCOS at various concentrations ranging from 250 to 4000 μg/mL for 24 h. The cell toxicity of HCOS on HUVECs and RAW 264.7 cells viability were then evaluated by the MTT assay performed as above. Cell viability was calculated as (absorbance value of sample/absorbance value of control) × 100%.

The hemolytic activity of HCOS was assessed according to a previously reported method (67) with some modifications. Briefly, 100 μL of a healthy rabbit red cell suspension in 96-well microtiter plates was mixed with 100 μL of a solution containing different concentrations of HCOS, and final HCOS concentrations of 250 to 4000 μg/mL were applied. After 1 h of incubation at 37 °C, the supernatants were collected, and the absorbance at 405 nm was measured using a TECAN Infinite M200 PRO multifunction microplate reader (TECAN, Grodig, Austria). Additionally, rabbit red cells incubated with PBS served as a negative control group, and those incubated with distilled water served as a positive control (serving as 100% hemolysis).

### Murine model of disseminated *candidiasis*

The sample size was selected based on the preliminary infection trial (n = 5 for mice models). Mice were relocated randomly to treatment or control cages. For the mouse disseminated Candidiasis model, Kunming female mice (n = 5 per group) were infected with a dose of 2.5 × 10^6^ *C. albicans* cells suspension by intravenous injection via the tail vein. Mice were treated 2 h post-infection with a specified intraperitoneal administration of fluconazole (0.1mg/kg, 5mg/kg) and oral gavage HCOS (10mg/kg, 50mg/kg) alone or the combination of fluconazole with HCOS (0.1mg/kg + 50mg/kg, 0.1mg/kg +10mg/kg), 0.9% NaCl as control. The animals were weighed every day, and after 7 days of treatment, the mice were euthanized, and one kidney was removed aseptically, placed in PBS, and then homogenized via bead beating. Serial dilutions of the homogenized kidneys were plated on YPD agar for the enumeration of fungal colonies. CFU values in kidneys were expressed as CFU/g of tissue, then transformed into log_10_ units. The differences between groups were analyzed by analysis of variance. Another kidney was fixed in 4% paraformaldehyde, embedded in paraffin wax, and sectioned longitudinally. Specimens were stained with periodic acid-Schiff (PAS) for the assessment of fungal invasion according to previous publication (68).

### Statistical analysis

Graphical evaluations were performed with GraphPad Prism v5.0 (GraphPad Software Inc., San Diego, CA). Analysis of variance (ANOVA) was used to evaluate significant differences. Data are presented as means ± SD. A two-tailed Student’s t-test was performed to compare two groups and one-way ANOVA for multiple group analysis. The P-value < 0.05 or 0.01 was considered as significant. All data were analyzed using SPSS 19.0 software (SPSS, USA).

## ACKNOWLEDGMENTS

We are grateful to Xixia. Li, and Xueke Tan for helping with sample preparation and taking TEM images at the Center for Biological Imaging (CBI), Institute of Biophysics, Chinese Academy of Science. We thank Prof. Huang for generously providing Phr1 mutant strains and thank Prof. Guan for providing GPI enzyme expression plasmid.

## FUNDING

This research was funded by National Key R&D Program of China grant number (2018YFC0311300) and National Natural Science Fund, China grant number (31670809).

## AUTHOR CONTRIBUTIONS

Conceptualization, Z.A.W. and Y.D.; methodology, Z.A.W., S.J., J.W., C.Z., L. R., X.G. and X.Y.; data acquisition, R.L., L.Z. and J.Z.; data interpretation, R.L.; software, D.L; visualization, Y.Z; writing-original draft preparation, R.L.; writing-review and editing, Y.Y and Z.A.W.; funding acquisition, Z.A.W. and Y.D. All authors have read and agreed to the published version of the manuscript.

## CONFLICTS OF INTEREST

The authors declare no conflict of interest.

## SUPPLEMENTAL MATERIAL

**Fig. S1.**
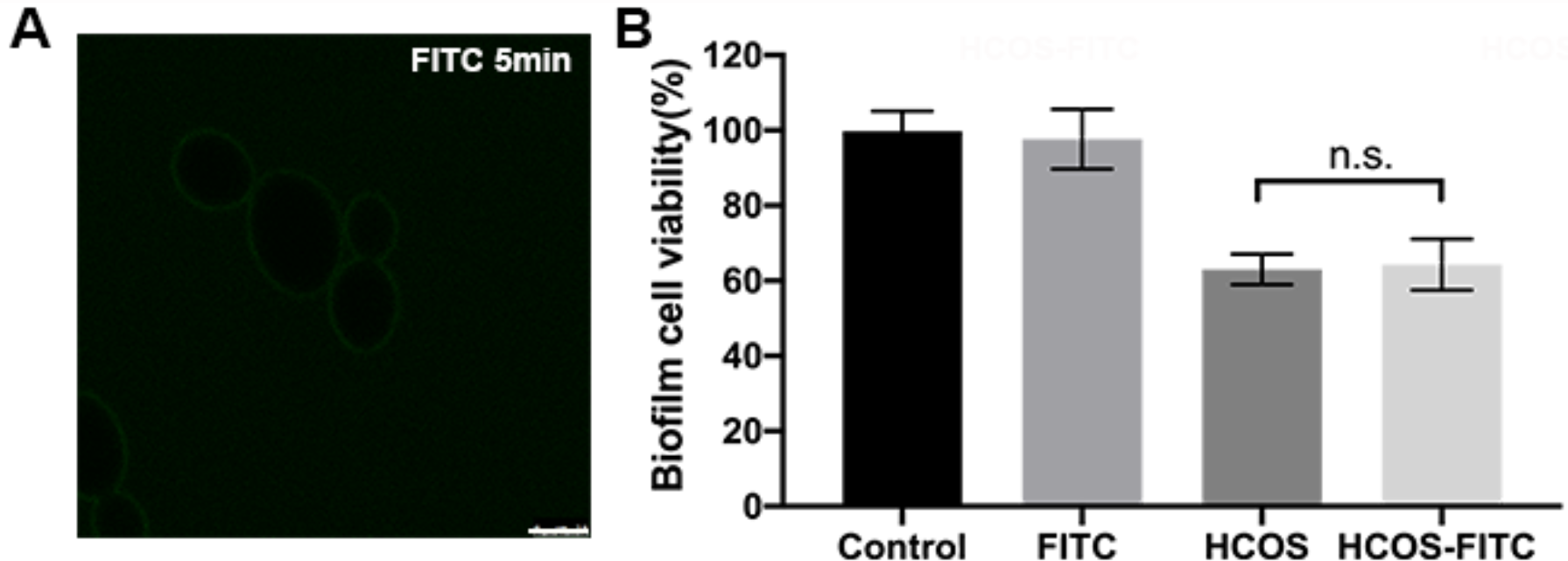
The antifungal activity of HCOS was not affected by FITC labeling. (A)The fluorescence microscope image of planktonic *C. albicans* treated with 500 μg/mL FITC for 5 min. Scale bar, 10 μm. (B) Mature *C. albicans* biofilm were treated with FITC, HCOS and HCOS-FITC at concentration of 500 μg/mL for 24 h, the biofilm cell viability was measured by XTT assay and normalized to the control. Data represented as means ± SD (n=6). n.s., non-significant.

**Fig. S2.**
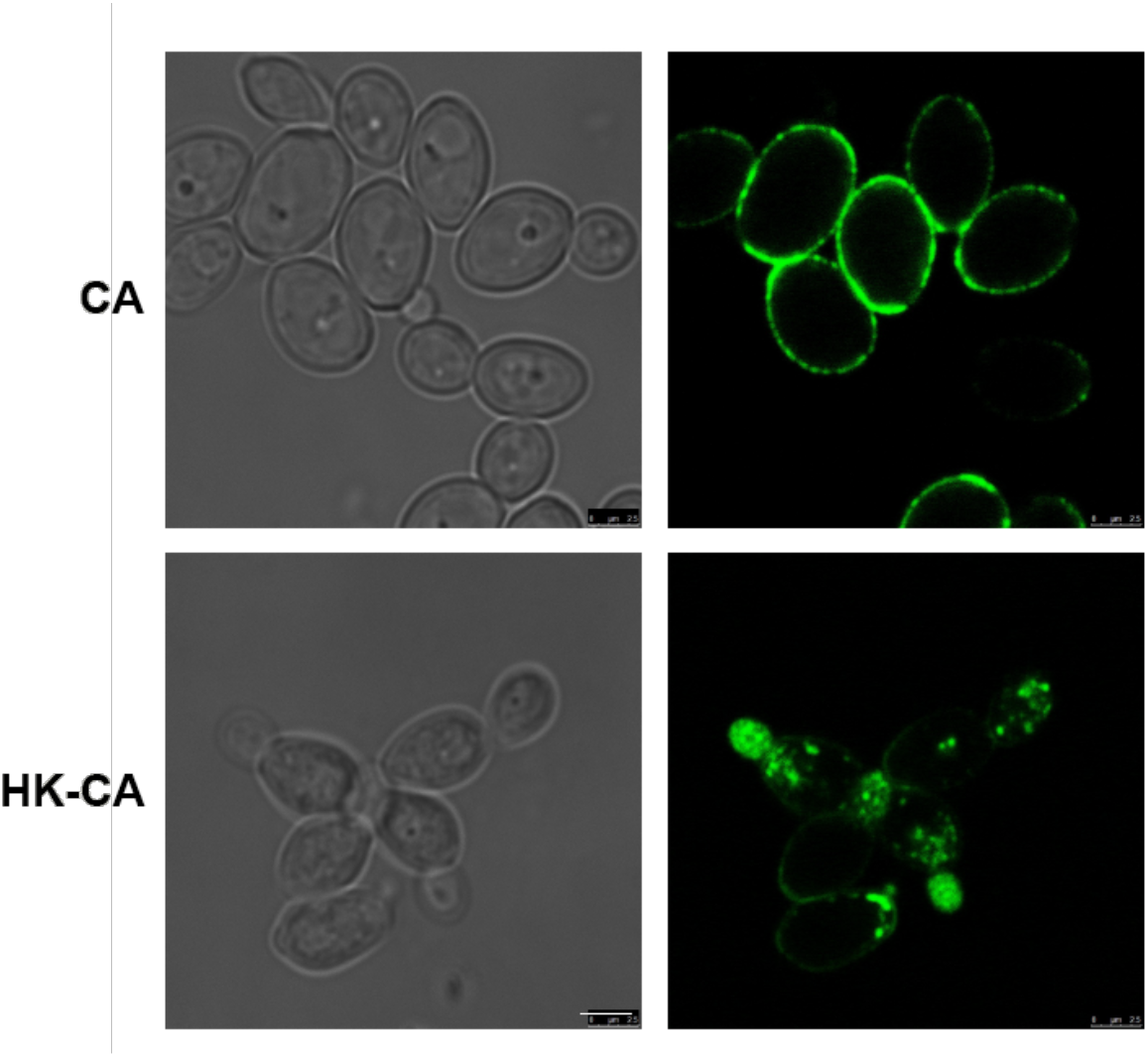
Fluorescence localization of HCOS-FITC on *Candida albicans* (CA) *and* heat-killed *Candida albicans* (HK-CA). Planktonic *C. albicans* and heat-killed *C. albicans* were incubated with 500 μg/mL HCOS-FITC for 1h at 30°C, 5 μL samples were transferred onto a slide and observed under fluorescence microscopy. scale bar, 10 μm.

**Fig. S3.**
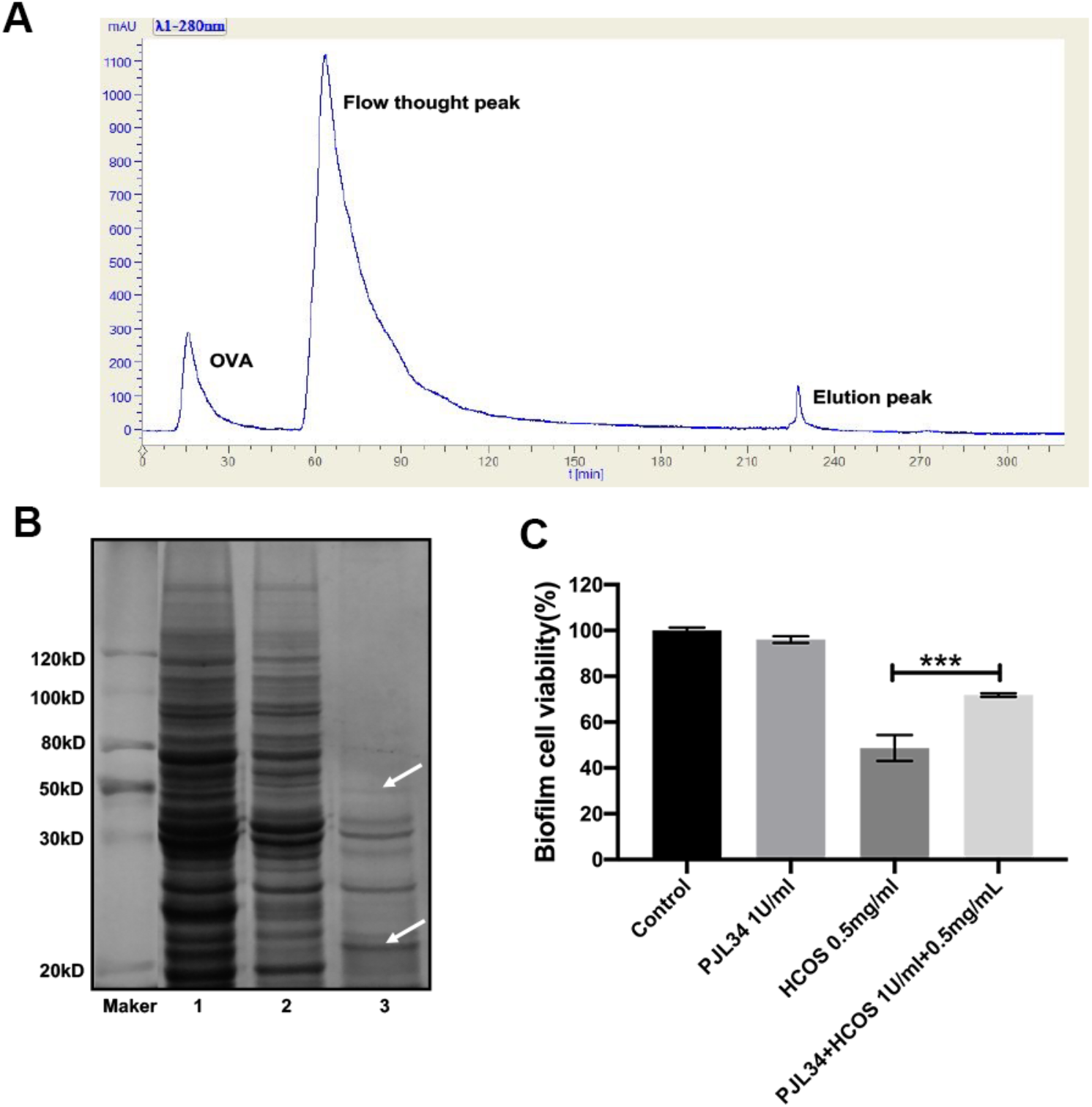
Purification and identification of HCOS-binding proteins in the *C. albicans* cell wall. (A) The process of isolating putative HCOS binding proteins using HCOS-Agrose-4FF chromatography column. (B) the SDS-PAGE analysis of the HCOS binding proteins. Coomassie blue staining of the 12% SDS-PAGE gel loaded with samples from the HCOS-Agarose-4FF affinity column. Line 1, the loaded sample; Line 2, flow-through; Line 3, the sample eluted from the HCOS-Agarose-4FF affinity column; Arrow indicated the potential HCOS-binding proteins. (C) The effects on C. albicans biofilm mass of HCOS and PJL34 (a β-(1,3)-glucanse from *Fibrobacter Succinogens*) alone or in combination for 24 h, the biofilm cell viability was determined by XTT assay and normalized to the control. Data are represented as means ± SD (n=6).

**Fig. S4.**
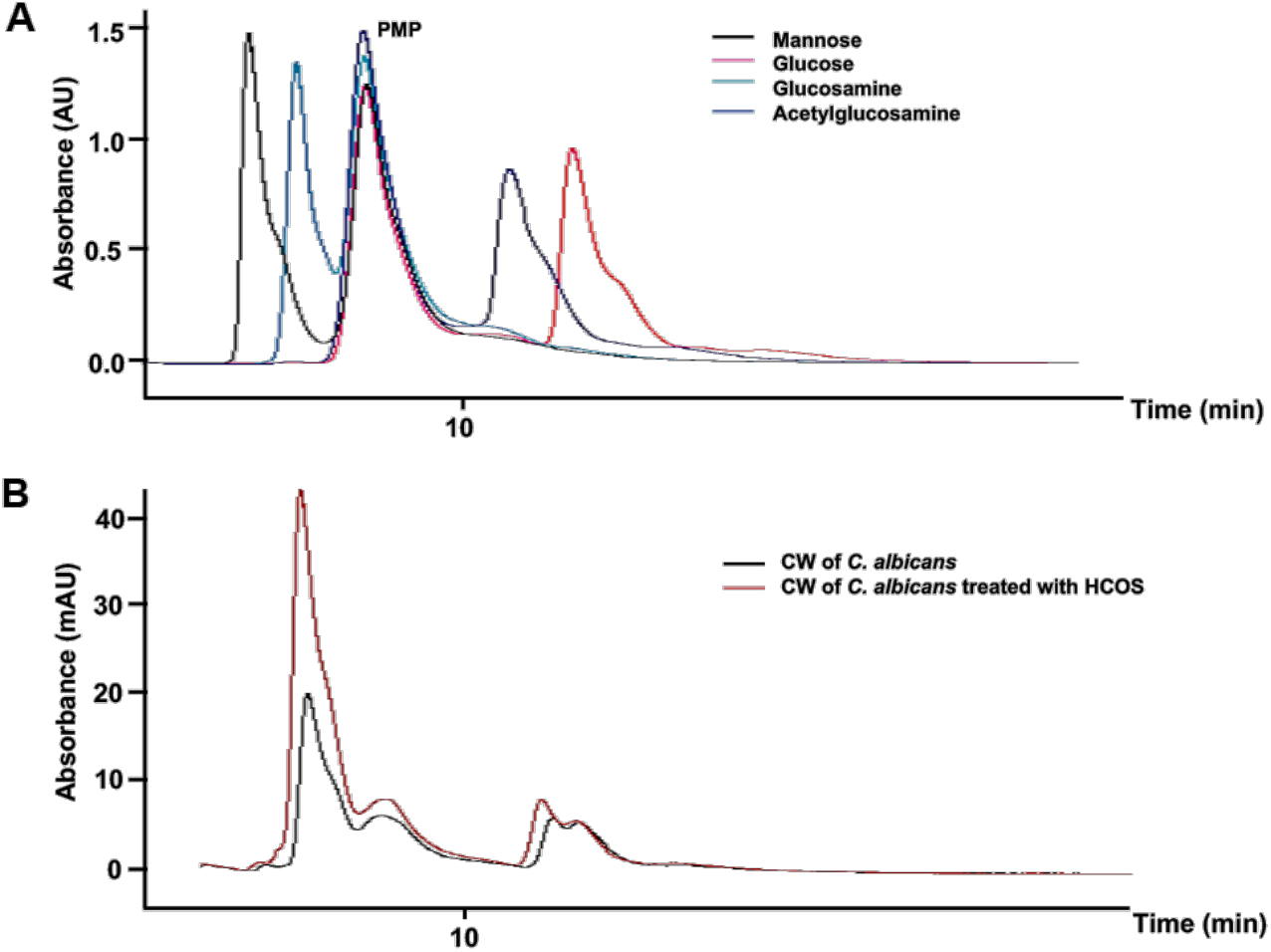
Degree of deacetylation of the cell wall chitin by HPLC analysis. (A) Four standard monosaccharides including mannose, glucose, glucosamine and acetylglucosamine were used as the references after being derivatized by 1-phenyl-3-methyl-5-pyrazolone (PMP). (B) HPLC analysis of TFAA hydrolyzed C. albicans cell wall, treated with or without HCOS (500 μg/mL), after being derivatized by PMP. The peak area of glucosamine of cell wall were 32.4% (control group) and 41% (CW treated with HCOS), respectively.

**Fig. S5.**
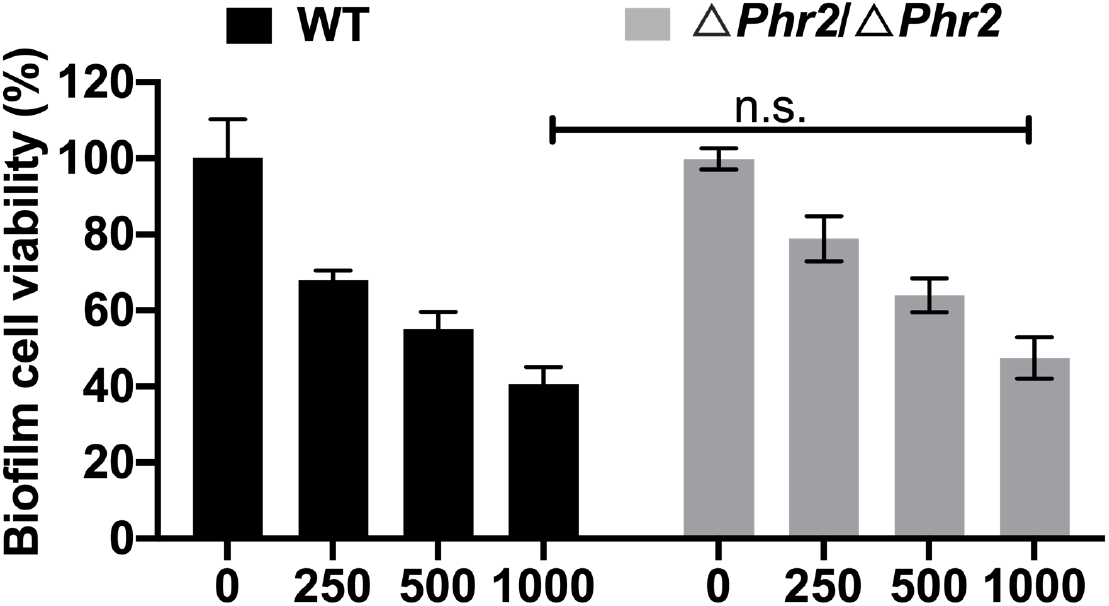
Susceptibility of a wild type C. albicans strain (GH1013) and a *Phr2* null mutant (Δ*Phr2*/Δ*Phr2*) to HCOS. The biofilm cell viability was determined by XTT assay and normalized to the control. Data represented as means ± SD (n=6).

**Fig. S6.**
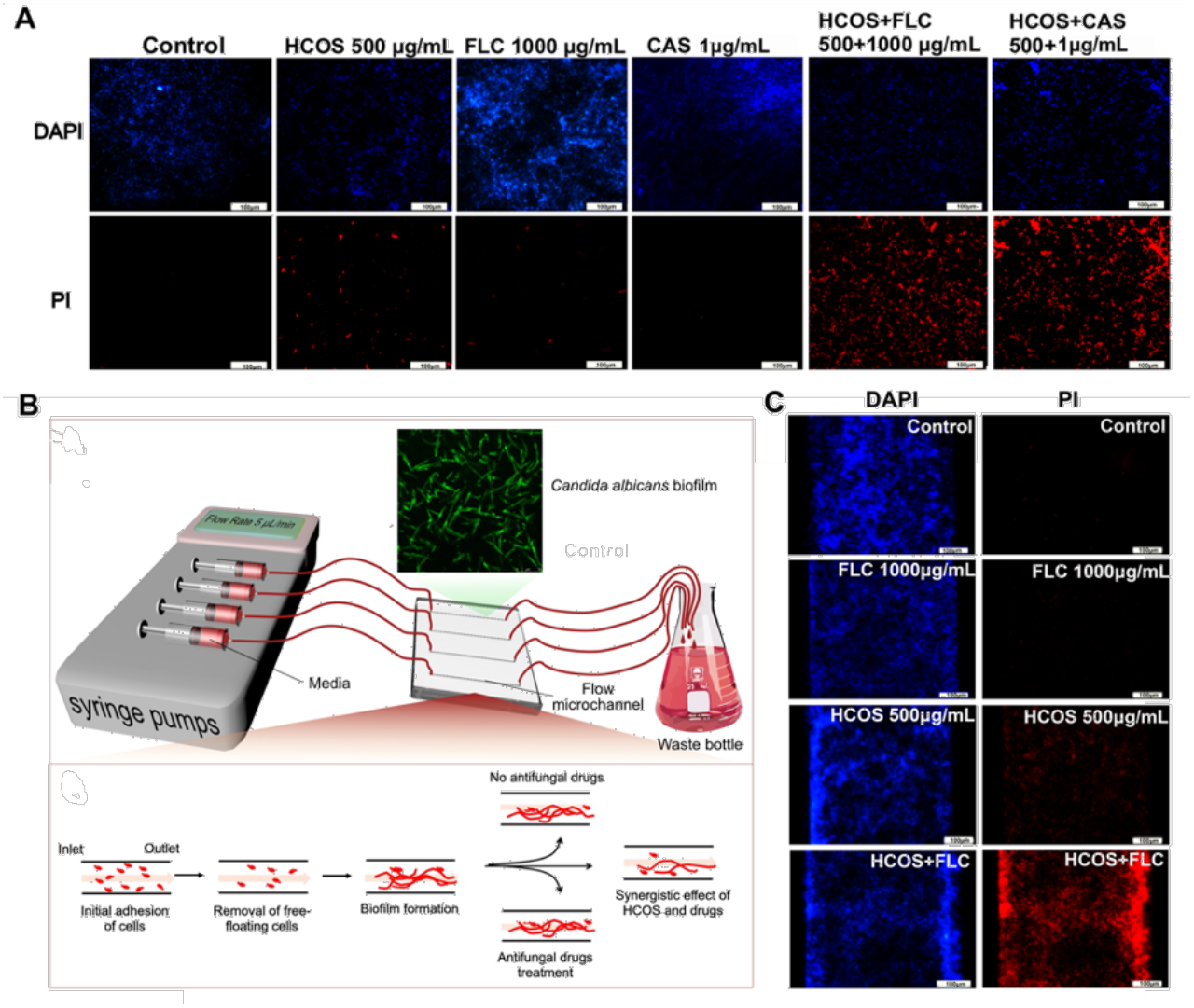
The synergistic antifungal effects of HCOS combined with fluconazole against *Candida albicans* biofilm. (A) Mature *C. albicans* biofilm formed on coverslip were treated with HCOS (500 μg/mL), FLC (1000 μg/mL), CAS (1μg/mL) and their mixture HCOS+FLC (500+1000 μg/mL), HCOS+CAS (500+1 μg/mL) for 24 h. Then the biofilm stained with 1μg/mL DAPI (blue) and 1μg/m PI (red) which represented total cells and dead cells, respectively. The samples were observed by fluorescence microscope. Scale bar, 100 μm. (B) Schematic diagram of the overall experiment of the biofilm microfluidic model. (C) *C. albicans* biofilms formed in a microfluidic chip and treated with fluconazole (1000 μg/mL), HCOS (500 μg/mL) or their combination with a flow rate of 5 μL/min for 2 h. The biofilm in microchannels was stained with 1μg/mL DAPI and 1μg/m PI and observed by fluorescence microscope. Scale bar, 100 μm.

**Table S1. The HCOS binding proteins candidates.**

**Table S2. Combined drug effects against planktonic *Candida* evaluated by the FICI model.**

MIC, the lowest concentration of an antimicrobial agent that inhibits the visible growth of microorganisms.

MIC_drug_, the MIC of antifungal agent when used alone.

MIC_HCOS_, the MIC of HCOS when used alone.

C_drug_, the MIC of antifungal agent when used in combination with HCOS.

C_HCOS_, the MIC of HCOS when used in combination with antifungal agent SYN, synergism.

**Table S3. Primers used in this study.**

## Notes

### Competing Interest Statement

The authors have declared no competing interest.

